# Coupling remote sensing and eDNA to monitor environmental impact: A pilot to quantify the environmental benefits of sustainable agriculture in the Brazilian Amazon

**DOI:** 10.1101/2023.07.19.549776

**Authors:** Karen Dyson, Andréa P. Nicolau, Karis Tenneson, Wendy Francesconi, Amy Daniels, Giulia Andrich, Bernardo Caldas, Silvia Castaño, Nathanael de Campos, John Dilger, Vinicius Guidotti, Iara Jaques, Ian M. McCullough, Allan D. McDevitt, Luis Molina, Dawn M. Nekorchuk, Tom Newberry, Cristiano Lima Pereira, Jorge Perez, Teal Richards-Dimitrie, Ovidio Rivera, Beatriz Rodriguez, Naiara Sales, Jhon Tello, Crystal Wespestad, Brian Zutta, David Saah

## Abstract

Monitoring is essential to ensure that environmental goals are being achieved, including those of sustainable agriculture. Growing interest in environmental monitoring provides an opportunity to improve monitoring practices. Approaches that directly monitor land cover change and biodiversity annually by coupling the wall-to-wall coverage from remote sensing and the site-specific community composition from environmental DNA (eDNA) can provide timely, relevant results for parties interested in the success of sustainable agricultural practices. To ensure that the measured impacts are due to the environmental projects and not exogenous factors, sites where projects have been implemented should be benchmarked against counterfactuals (no project) and control (natural habitat) sites. Results can then be used to calculate diverse sets of indicators customized to monitor different projects. Here, we report on our experience developing and applying one such approach to assess the impact of shaded cocoa projects implemented by the Instituto de Manejo e Certificação Florestal e Agrícola (IMAFLORA) near São Félix do Xingu, in Pará, Brazil. We used the Continuous Degradation Detection (CODED) and LandTrendr algorithms to create a remote sensing-based assessment of forest disturbance and regeneration, estimate carbon sequestration, and changes in essential habitats. We coupled these remote sensing methods with eDNA analyses using arthropod-targeted primers by collecting soil samples from intervention and counterfactual pasture field sites and a control secondary forest. We used a custom set of indicators from the pilot application of a coupled monitoring framework called TerraBio. Our results suggest that, due to IMAFLORA’s shaded cocoa projects, over 400 acres were restored in the intervention area and the community composition of arthropods in shaded cocoa is closer to second-growth forests than that of pastures. In reviewing the coupled approach, we found multiple aspects worked well, and we conclude by presenting multiple lessons learned.

## INTRODUCTION

Sustainable agriculture projects, which both generate income and contribute to environmental conservation, are important to address biodiversity loss, climate change, and to improve living conditions [1–3]. In Brazil, sustainable agriculture approaches have been developed in response to the simultaneous pressures of combatting ongoing habitat and biodiversity loss caused by agricultural expansion and food security issues [4–10]. Such practices include agroforestry, sustainability certifications, and the promotion of non-timber forest products.

Environmental monitoring is essential to ensure that the promised impacts of sustainable agriculture are being achieved [11] and a growing interest in environmental monitoring provides an opportunity to improve monitoring practices [1,12–16]. To improve transparency and enhance credibility, organizations need accurate, timely, and easily digestible information collected using robust methods informed by the best available science. Because the environmental impacts of projects or interventions can take many years to become evident, any monitoring system must be replicable and comparable across time. Monitoring systems should examine multiple scales to account for both individual farm and landscape-scale habitat loss or fragmentation, habitat diversity, and connectivity [7, 17–18]. Similarly, to ensure that the measured impacts are due to the project and not exogenous factors, sites where projects have been implemented should be benchmarked against counterfactuals (no projects) and control (natural habitat) sites. Indicators, or predefined metrics for assessing ecosystem services, ecosystem health and biodiversity, can be used to assess relative performance rapidly and can be designed with communication to both experts and non-experts in mind.

Using remote sensing and environmental DNA (eDNA) approaches, systematic, broad-scale, multi-year monitoring efforts are financially and operationally feasible. Traditional approaches to monitoring forests and biodiversity, such as forest surveys and biodiversity transects, are expensive and require significant methodological and taxonomic expertise, particularly in megadiverse regions [19]. In contrast, products derived from remote sensing facilitate a substantial reduction in monitoring costs and simultaneously increase the timeliness of information needed to inform management [20–22]. Remote sensing uses satellite and aircraft imagery and statistical approaches to detect and monitor the Earth. Existing remote sensing approaches for monitoring biodiversity allow for the evaluation of ecosystem structure and ecosystem function but are not yet extensively used in biodiversity assessment, monitoring, or conservation [23].

Similarly, eDNA monitoring has speed and cost advantages over traditional methods; thus, it is rapidly becoming a preferred method to monitor biodiversity, including in agricultural systems [24–27]. eDNA refers to genetic material obtained directly from environmental samples, such as water, soil, or air, without capturing or observing the organisms themselves [28] and coupled with the metabarcoding approach allows for the simultaneous identification of multiple species from a single sample [29]. eDNA sampling approaches can monitor entire taxonomic groups at multiple spatial scales depending on whether eDNA is collected from leaves, soil, animal waste, water, or air [30–35].

Coupling the two technologies allows for both wall-to-wall coverage from remote sensing and the site-specific community composition from eDNA (Figure 1). Together, these methods can be used to calculate diverse sets of indicators for monitoring sustainable agriculture projects and other sustainable interventions. These indicators can include but are not limited to measures of ecosystem function such as landscape scale and site scale habitat loss and conservation, ecosystem structure, carbon sequestration through revegetation, species richness, community composition, and relative changes in community composition over time. The approaches are cost effective, and the results provide a complete picture of activities occurring across the full project site.

**Figure 1:**
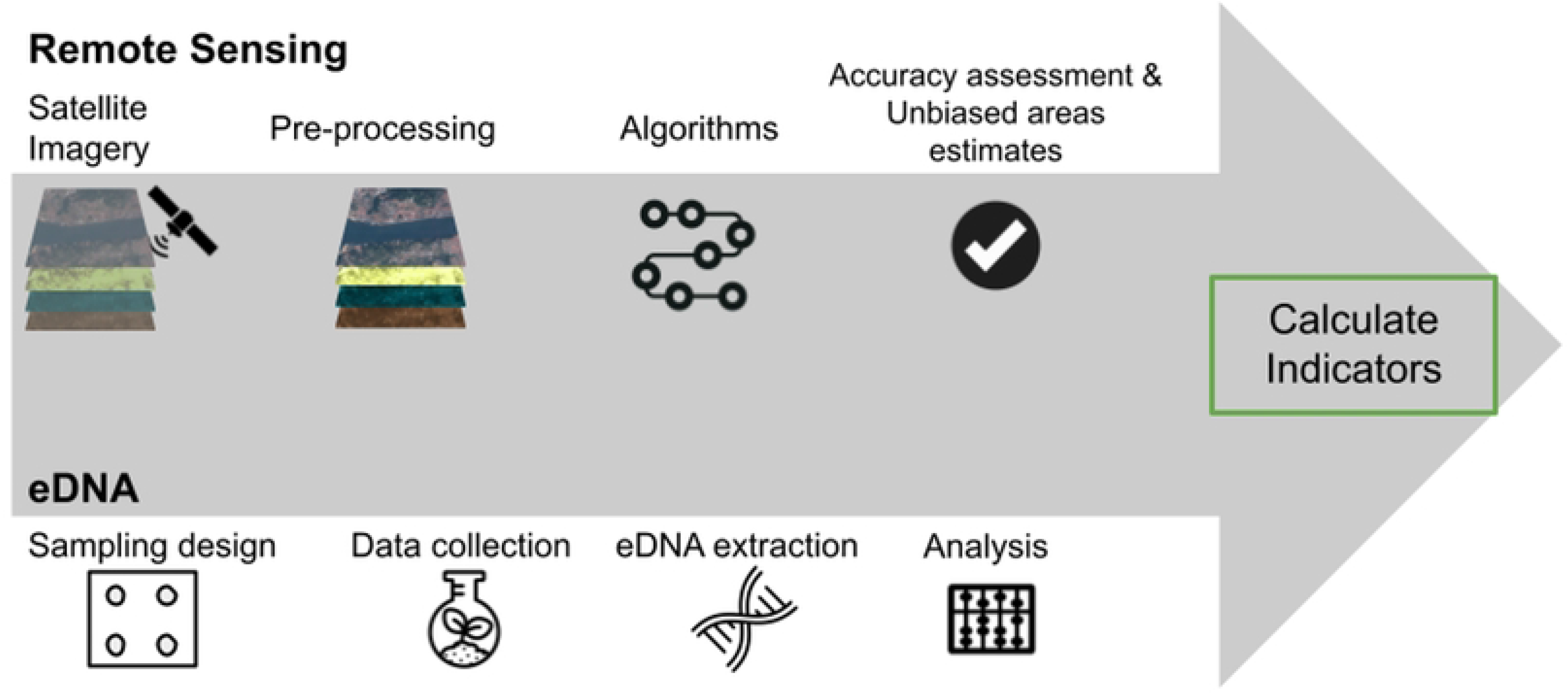
An overview of the proposed coupled approach to environmental monitoring. In the remote sensing component, we identify and pre-process key satellite imagery. Next, we input data into change algorithms or other models. Then, we assess the accuracy of the produced maps. In the eDNA component, we design the sampling approach by identifying locations to sample. Next, we visit sites to collect eDNA data. Then, the soil samples go through the eDNA extraction and biodiversity analysis processes. Finally, we calculate indicators from the map outputs and biodiversity results.

Here, we report on the pilot application of this approach to assess the effect of one of the sustainable agriculture initiatives of *Florestas de Valor*, created by the Instituto de Manejo e Certificação Florestal e Agrícola (IMAFLORA). *Florestas de Valor* consists of multiple initiatives including agroforestry and collection of non-timber forest products [36]. We applied the coupled monitoring approach to shaded cocoa projects implemented by IMAFLORA near São Félix do Xingu, in Pará, Brazil. Shaded cocoa in Brazil is currently being supported as an alternative to unshaded cocoa and low-yield pasturelands used for cattle ranching [37–38]. Key research questions included: 1) have the shaded cocoa projects contributed to conservation in the Brazilian Amazon? and 2) what impacts does shaded cocoa have on community structure and forest landscape patterns? We tested the relative effects of these management activities on biodiversity conservation and compared the results with counterfactual “business as usual” pastures and control second-growth forests to control for outside factors.

This implementation of coupling remote sensing and eDNA for biodiversity monitoring provided an opportunity to test multiple approaches and improve the methodology. Key areas of testing included our remote sensing mapping approach, including algorithm selection, sampling design and our data collection approach, and indicator choices. These tests contributed to lessons learned that will greatly improve coupled biodiversity monitoring methods moving forward.

## MATERIALS AND METHODS

### STUDY AREA

The study area is in the Xingu River basin, near the city of São Félix do Xingu, in the state of Pará, Brazil (06°38′30″South and 51°58′32″West). The region is warm moist equatorial, with dry months from May to October. The average annual rainfall is 2,041 mm. The average annual temperature is 25 °C, with minimum and maximum temperatures of 20 °C and 30 °C, respectively [39]. Native tree species in the region include *Attalea speciosa* (Mart.) and *Cedrela odorata* L., for example [40]. Primary forest cover was almost entirely removed from the area in the 1960s for agriculture and was slowly replaced by secondary vegetation, including forest. Subsequently, farmers cleared secondary forests for cattle grazing [41–42].

More recently, shade-grown cacao (*Theobroma cacao* L.) has been promoted in the region as a sustainable alternative [42–44]. Shaded cocoa is thought to reduce agricultural inputs, disease susceptibility, and drought susceptibility as well as increase food security and environmental benefits [43–44]. There are two phases to cultivating shade trees in cocoa agroforestry systems. First, specific shade trees, including banana and papaya, are cultivated while native regeneration occurs, and second, these cultivated trees are thinned, and native shade trees become dominant [44]. During the first phase, annual crops like cassava and maize are also grown, with cassava chosen to increase the nitrogen content in the soil [44]. Native shade trees include *Apuleia leiocarpa* (Vogel) J.F. Macbr., *Bagassa guianensis* Aubl., *Pouteria macrophylla* (Lam.) Eyma, *Erythrina verna* Vell., *Pouteria pariry* (Ducke) Baehni, *Chrysophyllum cuneifolium* (Rudge) A. DC., *Perebea guianensis* Aubl., *Spondias mombin* L., *Colubrina glandulosa* Perkins, *Cenostigma tocantinum* Ducke, *Annona mucosa* Jacq., *Handroanthus serratifolius* (Vahl) S.Grose, *Inga edulis* Mart. and *Samanea tubulosa* (Benth.) Barneby and J. W. Grimes [44].

IMAFLORA is a nonprofit partner in the SERVIR Amazonia consortium based in Brazil. IMAFLORA maintains a database of 150 farms participating in multiple agricultural practices and has worked with a subset of these farmers to implement shaded cocoa practices over the past 20 years [45, Figure 2]. Within this context, our study boundary encapsulated the farms partnering with IMAFLORA.

**Figure 2:**
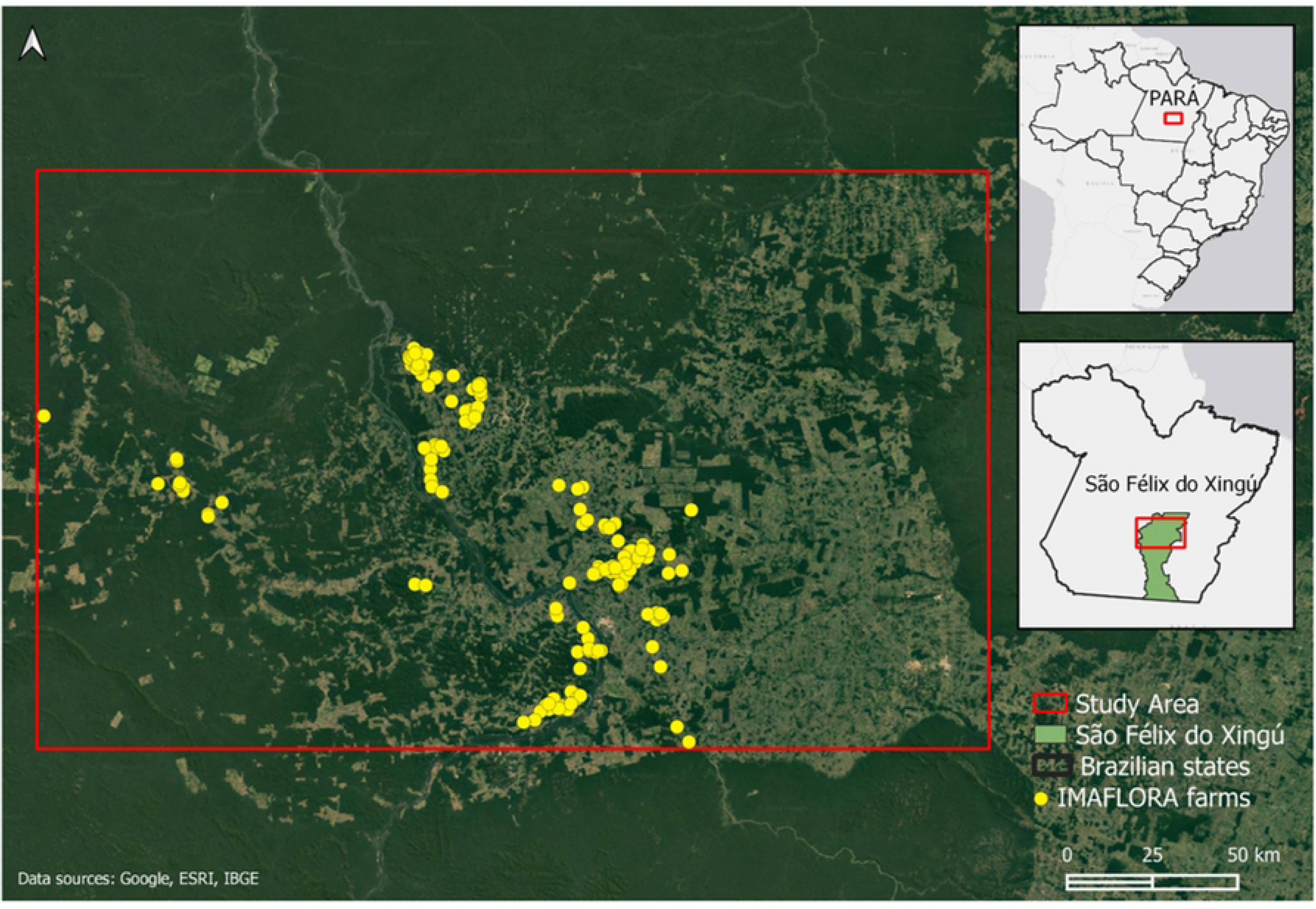
Study area in the state of Pará, Brazil. Yellow dots represent the farms that have partnered with IMAFLORA.

### REMOTE SENSING METHODS

#### DISTURBANCE MAPPING

Forest disturbances were mapped using a combination of two pixel-based methods: the Continuous Degradation Detection (CODED) algorithm [46–47], and the Landsat-based detection of Trends in Disturbance and Recovery (LandTrendr) algorithm [48]. Both algorithms utilize Landsat collections in a time series approach (30 m spatial resolution). Here, disturbances are separated into deforestation and forest degradation. We consider deforestation a permanent conversion of forested land to non-forested land [49], and degradation a process that does not lead to a categorical land cover change but shows the loss in tree cover canopy [50–51]. The changes were mapped for the 5-year period 2010-2015, when most of the interventions occurred. The results of the two algorithms were combined and evaluated for accuracy.

CODED is a freely available tool on Google Earth Engine (GEE), an online planetary-scale computing platform for remote sensing and satellite imagery analysis [52]. CODED uses all the Landsat imagery available from Landsat collections 5, 7, and 8 to perform a subpixel spectral mixture analysis (SMA), analyzing time series changes in the Normalized Degradation Fraction Index (NDFI; 30 m resolution; 46, 50). The spectral index of choice was the NDFI to be in accordance with CODED and also since previous studies have shown that NDFI is more sensitive to disturbances in tropical forests compared to the commonly used Normalized Difference Vegetation Index (NDVI), which tends to show higher variability [46–47, 53]. We used 3,000 training points labeled as forest or non-forest and tested chi-squared values of both 0.9 and 0.99, which controls the width of the change-detection moving time window [51]. We defined the required number of sequential out-of-range NDFI values to flag an event as four times. In post-processing, we required the magnitude of these flagged disturbances to be above 0.4 to be considered severe enough to include in the final map. In this manner CODED can detect low-severity disturbances, which is often characteristic of the more difficult-to-detect forest degradation events, as opposed to forest loss. The final CODED output is a map with 30 m pixels labeled as non-forest, stable forest, deforestation, or degradation.

Next, we used the LandTrendr algorithm, also freely available and hosted on GEE. LandTrendr is a collection of algorithms used to detect land cover change through time series analysis of Landsat imagery. Specifically, LandTrendr constructs an image collection by creating medoid composites of Landsat 5, 7, and 8 images, resulting in one image per year. The medoid image compositing approach compares each pixel’s spectral band values to the median spectral values of those bands across all images within the date-constrained collection for a given year. The pixel with spectral values closest to the median value, determined by Euclidean spectral distance, is then selected [48]. This tool aims to filter out inter-annual noise in spectral signals and generate trajectory-based time series estimates and accomplishes this through simplifying multi-year spectral trajectories into several straight-line segments that capture the progressing changes of the signal [48]. The algorithm was parameterized to estimate the “greatest” disturbance, and specific parameters are mostly in accordance with previous research [54, 55]. We further classified the disturbances as degradation and deforestation using MapBiomas classification in 2015 [56]. Disturbances classified as forest by MapBiomas in 2015 were considered degradation, whereas disturbances classified as non-forest were classified as deforestation.

Lastly, the final disturbance map was generated by overlapping the CODED and LandTrendr maps and using a rule-based assignment, where pixels with classification disagreements were reclassified as: degradation if at least one of the outputs was classified as degradation, deforestation if none of the outputs were classified as degradation and at least one classified as deforestation, and, consequently, forest if both outputs were classified as forest. We used this system because degradation was underestimated in previous studies [51, 54, 57].

#### REGENERATION MAPPING

Similarly to the disturbance mapping, we used LandTrendr to map forest regeneration following previous research [55, 58]. LandTrendr was set up to detect upward trends in the spectral signature of forested areas. The same NDFI index was used, and the parameterization was set to estimate the “greatest” gain and specific parameters adjusted as an attempt to capture short-time regeneration given the 5-year period.

Based on maps by the Brazilian Annual Land Use and Land Cover Mapping Project (MapBiomas), [59] developed maps of annual secondary forest extent, age, increment, and loss within Brazil for 1986-2019 [56]. Using MapBiomas data, secondary forest growth is present when a pixel with an anthropic cover classification (e.g., pasture or agriculture) becomes a forest cover pixel (excluding mangroves and forest plantations) the following year [59]. We utilized the same methodology to generate a map of secondary forests, here also considered as regeneration, from 2010 to 2015. These maps were also Landsat-based with a 30 m spatial resolution.

The final regeneration map was a combination of the LandTrendr output with the output from [59] methodology where a pixel is mapped as “regeneration” if at least one of the outputs was mapped as “regeneration”, in an “all-inclusive” approach. This way, we could assess this approach’s accuracy and inform future TerraBio work.

#### ACCURACY ASSESSMENT OF MAPPING PRODUCTS

An independent validation effort was conducted leveraging high- and medium-resolution optical imagery (Planet NICFI mosaics, Google Earth Pro basemaps, Sentinel-2, and Landsat Collections) and ancillary datasets (MapBiomas products, the Global Forest Canopy Height 2019, and NDFI time series) to assess classes on the ground [60–65]. Visual interpretation was done in Collect Earth Online (CEO), a free and open-source web-based tool that facilitates data collection and validation [66–67]. The interpreter utilized a decision tree approach for classifying the validation samples.

For the validation of the disturbance and regeneration maps, the sample points were 30 by 30 meters square, mimicking the map output pixel sizes. The validation of both change maps (disturbance and regeneration) was done using the same set of points. We also used a simple semi-random sampling design with proportional class distribution where we extracted 600 points from both maps, therefore, 100 points for each class: degradation, stable non-forest, stable forest, deforestation, regeneration, different events (Table S1). A “different events” category was used to capture locations where the mapping methods determined a different class for the same location. This class can include regeneration events on non-forest areas followed by degradation or deforestation or disturbance events, most likely deforestation, followed by regeneration. The accuracy metrics (overall, user, and producer accuracies) and unbiased area estimates for each class were calculated through the ratio estimator approach [68] for when the strata are different from the map classes since we used the same sample points for disturbance map and regeneration map.

### eDNA METHODS

#### eDNA FIELD DATA COLLECTION

We defined our sampling frame as the 150 farms partnering with IMAFLORA. Within this sampling frame, we verified which farms included our project sites, or farms with mature shaded cocoa, by checking average vegetation height of cocoa fields (minimum average canopy height of 7.5m) and verifying that fields contained greater than 25% canopy cover in CEO [65]. We identified 49 farms containing mature shaded cocoa.

Our sampling design was chosen to reduce the influence of exogenous variables that influence biodiversity to maximize our statistical power in detecting differences between project and counterfactual sites. Key exogenous variables include the amount and configuration of forest both within the farm boundaries and in a 250m buffer [69–73]. To represent these exogenous influences, we selected a suite of landscape ecology metrics that best captured landscape variance using principal component analysis (PCA; 74-75). Metrics used for clustering included within the farm boundaries: total forest area, number of forest patches, forest percentage of farm, forest contiguity, and forest aggregation index; and within the 250m buffer included: total forest area, forest percentage of landscape, and forest aggregation index [76]. We used these landscape metrics to assign each farm to a cluster using Ward’s hierarchical clustering with the Euclidean distance matrix [77].

Following cluster assignment for each farm, we used stratified random sampling to select five project sites and five counterfactual sites with the same proportion of each cluster in each group of sites. We also used the same process to select two back-up sites for each group. We avoided excluding clusters since that would have changed the sampling frame. In addition, experts from IMAFLORA identified five forest sites that represented the oldest known second growth forest areas in the study area. However, the field team experienced difficulties accessing sites, weather delays, and COVID delays. This led to only one second growth forest being sampled, reduced numbers of plots sampled in one shaded cocoa field and two pastures, and the use of both shaded cocoa backup sites and one pasture backup site.

A total of 38 plots were sampled, with 18 plots from 5 shaded-cocoa sites, 17 from 5 pasture sites and 3 from one forest site. Within each site, we randomly placed either 3 or 4 50m x 50m plots depending on the field size, with at least 25m between plots. Each plot had four sub-samples, set back from the edge of the plot by 12.5m, to better sample the heterogeneity found within the plot. Approximately 30g of the topsoil (between 0-5 cm deep) was collected for each sub-sample, making sure to avoid non-soil matter including leaf litter, and sub-samples were pooled to represent one sample per plot. To avoid sample contamination and ensure consistency across samples, a protocol was followed during sampling that included the use of disposable sampling materials and gloves and samples were labeled and individually packed according to plots and sites to ensure no cross-contamination occurred (detail of methods see S2 File; 78). Following collection, soil samples were stored with silica desiccant bags (minimum of 10g per sample) protected from heat and sunlight to prevent DNA degradation and later transported to the laboratory facilities for further laboratory analysis.

#### eDNA EXTRACTION

Soil samples were pre-processed in a Department for Environment, Food & Rural Affairs (DEFRA) licensed laboratory facility. To avoid contamination, samples were handled in a pre-PCR laboratory, using disposable tools and gloves, following standard decontamination procedures (i.e., use of bleach to clean surfaces and equipment), and personnel wore disposable full-body suits when handling the samples.

The extraction method was conducted using 2 g of mixed soil (per analyzed plot) and following the Mu-DNA soil DNA extraction protocol described by [79]. Negative controls were included, comprising DNA extraction blanks containing only the required buffers.

Following DNA extractions, DNA amplification was conducted using three sets of primers targeting two partial mitochondrial genes. First, vertebrate specific primers were used targeting ∼106 bp of the 12S rRNA gene (80; forward primer 5’-TAGAACAGGCTCCTCTAG-3’ and reverse primer 5’-TTAGATACCCCACTATGC-3’). Second, to detect arthropods, DNA extracts were amplified using two primer sets targeting different short inserts of the mtDNA COI gene. The Zeale primer set [81] was used to amplify a ∼157 bp fragment, and the Gillet primers [82–84] were used to amplify a ∼133 bp section (Zeale: forward primer 5’-AGATATTGGAACWTTATATTTTATTTTTGG-3’ and reverse primer 5’-WACTAATCAATTWCCAAATCCTCC-3’; Gillet: forward primer 5’-CCATCTCATCCCTGCGTGTCTCCGACTCAGNNNNNNNATTCHACDAAYCAYAA RGAYATYGG-3’ and reverse primer 5’-CCTCTCTATGGGCAGTCGGTGATNNNNNNNACTATAAAARAAAATYTDAYAAA DGCRTG-3’).

PCR reactions consisted of 12.5 µl Master Mix, 7.5 µl molecular grade water, 2 µl of DNA template and 1 µl of the forward and reverse of each primer. The PCR conditions for the Riaz primer followed the [85] methodology, consisting of an incubation of 5 minutes at 95 °C, then 35 cycles for 15 seconds at 95 °C, 30 seconds at 57 °C, ending with 30 seconds at 72°C. PCR conditions of the Gillet and Zeale primers followed the protocols set by [86]. Gillet cycles included an initial 15-minute denaturation at 95 °C, then 10 cycles for 30 seconds at 94 °C, 45 seconds at 49 °C, 30 seconds at 72 °C, 30 cycles of 30 seconds at 95 °C, 45 seconds at 47 °C and 30 seconds at 72 °C, with a final extension following of 10 minutes at 72 °C. Zeale PCR conditions began with a 15-minute denaturation at 95 °C, then 40 cycles of 20 seconds each at 95°C, 30 seconds at 55 °C and 1 minute at 72 °C, with a final extension of 7 minutes at 72 °C. PCR cycles were authenticated by electrophoresis in a 1.2% agarose gel stained with GelRed. PCRs were run in triplicates, and the success of the reactions was determined by electrophoresis on a 1.5% agarose gel. Four PCR blanks were included in each library to account for putative contaminations arising in the amplification steps. In total, 44 samples were analyzed per library, 38 eDNA soil samples, two extraction blanks and four PCR blanks. A left-sided size selection was performed using 1.2× Agencourt AMPure XP (Beckman Coulter) and the KAPA HyperPrep kit (Roche) was used to construct the Illumina libraries using the dual-indexed adapters. Libraries were quantified using the NEBNext qPCR quantification kit (New England Biolabs) and pooled in equimolar concentrations. Two Illumina MiSeq sequencing runs were conducted, one MiSeq v2 Reagent Kit (2 × 150 bp paired-end reads) and one MiSeq v3 Reagent Kit (2 × 300 bp paired-end reads).

Bioinformatic steps were conducted as described in [86]. In brief, bioinformatic analysis used the OBITools 1.2.2 metabarcoding package [87]. Read quality was assessed using FastQC, Illumina paired end aligned paired-end reads, and ngsfilter demultiplexed samples and removed primers. The obigrep command performed size selection by eliminating artifacts and ambiguous reads. Vsearch [88] clustered unique sequences and removed chimeras using uchime-denovo [89]. Sumaclust clustered sequences into Molecular Operational Taxonomic Units (MOTUs) at thresholds of 0.95-0.98. Taxonomic assignment relied on Basic Local Alignment Search Tool (BLAST, specifically blastn) against Genbank, with a minimum of 90% alignment and >80% similarity [90]. Species-level assignment required ≥98% identity, while MOTUs at 95%-98% or with multiple species were assigned at the genus level. MOTUs between 93%-95% were assigned to the family level, and MOTUs between 90%-93% were assigned to the order level [86]. Sequences were retained when they could be identified at least to the Class level. A final filtering step was conducted, including the removal of putative contaminants, tag-jumping (MOTUs represented by less than 0.01% of the total reads were removed from each sample), and non-target taxa (e.g., Human DNA). Additionally, molecular operational taxonomic units (MOTUs) were retained when the total number of reads was over 50 [91]. MOTUs were considered compositional data and treated as such [92–94], except for when indicators called for species abundance measures where reads were used [95].

#### eDNA ANALYSIS

The four initial biodiversity indicators and four proposed indicators were calculated based on the resulting data. For indicators using key species, we defined key species for mammals as threatened native species and also excluded domesticated and invasive species from analysis more broadly [96]. We defined key species for arthropods as members of the Hymenoptera and Lepidoptera orders as important pollinator species, including for coffee crops [97–98]. These indicators require BLAST to match sequences to the Order level. As has been previously reported in other studies [e.g., 86], the Gillet primers were notably better at detecting members of Hymenoptera, and the Zeale primers were better at detecting members of Lepidoptera. Due to the limited overlap between the two primer sets, we combined the resulting datasets for further analysis. For determining if any MOTUs were associated with (indicative of) either cocoa fields or pasture and the ecological conditions found there [99–100] we used the multipatt function {indicspecies} with a custom wrapper [101–103].

For community-based indicators, we used all sequences identified by BLAST as Arthropods, requiring identification to the Phylum level. We first accounted for zeros in the dataset using zCompositions [104], then transformed the data using compositions [105]. We calculated Aitchison distance between sites and between treatments using the Euclidean distance matrix [26, 106–107]. We created PCA plots using the transformed compositional data [106–107].

Where needed, we used linear mixed models to account for the repeated sampling design to test the difference between projects and counterfactuals [108]. Significance tests were performed using ANOVA and Type II Sums of Squares [109]. All data analysis was conducted in R [110] and all code is available on GitHub (https://github.com/sig-gis/TerraBioPilot).

### CARBON CALCULATIONS

Carbon sequestration through revegetation estimates was calculated using modified methods based on the New IPCC Tier-1 Global Biomass Carbon Map for the Year 2000 [111]. Tier 1 carbon estimates are defined using a look-up table which has an associated carbon value based on Global Land Cover 2000 (GLC2000) cover type, ecofloristic zone, continental region, and frontier forest designation [112]. We assumed that the pre-regeneration GLC2000 land cover classification was the Cultivated and Managed land class. For regeneration areas, we used the Tropical Rainforest ecofloristic zone based on the UN FAO maps, modified with their 50% factor for disturbed vegetation categories [113]. Areas of carbon gain were determined based on areas of gain from the regeneration map product located in the shaded cocoa areas for the 150 farms.

### MORPHOLOGICAL SPATIAL PATTERN ANALYSIS (MSPA)

To calculate the number of hectares of essential habitat areas we used the Morphological Spatial Analysis (MSPA) which takes a binary image composed of the objects of interest (foreground area) and complementary data and divides it into morphological classes, or classes that describe the spatial arrangement of habitat across the landscape [114]. This is performed through a customized sequence of mathematical morphological operators targeted at the description of the geometry and connectivity of the image components. The MSPA segmentation results in 25 mutually exclusive feature classes which, when merged, exactly correspond to the initial foreground area. In this case, we used the binary image corresponding to loss areas from 2001 to 2019 of the Global Forest Change dataset [115] as input. The image output had 8 classes: patch forest, outer edge, inner edge, core forest, secondary degradation, secondary deforestation, primary degradation, and primary deforestation (for definitions, see S2 File)

### TERRABIO

To assess the shaded cocoa projects implemented by IMAFLORA, we conducted a pilot of TerraBio. TerraBio is a methodology framework developed to provide environmental assessment and accountability to private sector firms that (1) commercialize sustainable agriculture and forest products and/or, (2) invest in sustainable business models as profitable and conservation-driven development initiatives. TerraBio directly monitors the land cover change and biodiversity measures annually to provide timely and relevant results for investment funds, businesses, and other parties invested in the success of sustainable agricultural practices [116].

TerraBio uses a coupled approach to environmental monitoring, with the landscape component conducted with remote sensing and the biodiversity component conducted with eDNA, which work together to calculate a series of indicators. Intervention areas were shaded cocoa fields; our counterfactual areas or ‘business as usual’ areas were pasture fields, and our control areas were the areas of naturally regenerated second growth forest.

The initial set of indicators for the TerraBio pilot were derived from existing sources, including the Amazon Biodiversity Fund Brazil Key Performance Indicators (KPI) and the United States Agency for International Development/Brazil’s Partnership for the Conservation of Amazon Biodiversity standard indicators [116–118]. These indicators were: Number of hectares directly restored; Number of hectares indirectly conserved; Carbon sequestration through revegetation; Number of keystone/priority species; Change in abundance of keystone/priority species; Change in species richness; Change in biodiversity index; Number of hectares showing improved biodiversity; and Number of hectares of essential habitat area conserved (Table 1).

**Table 1:**
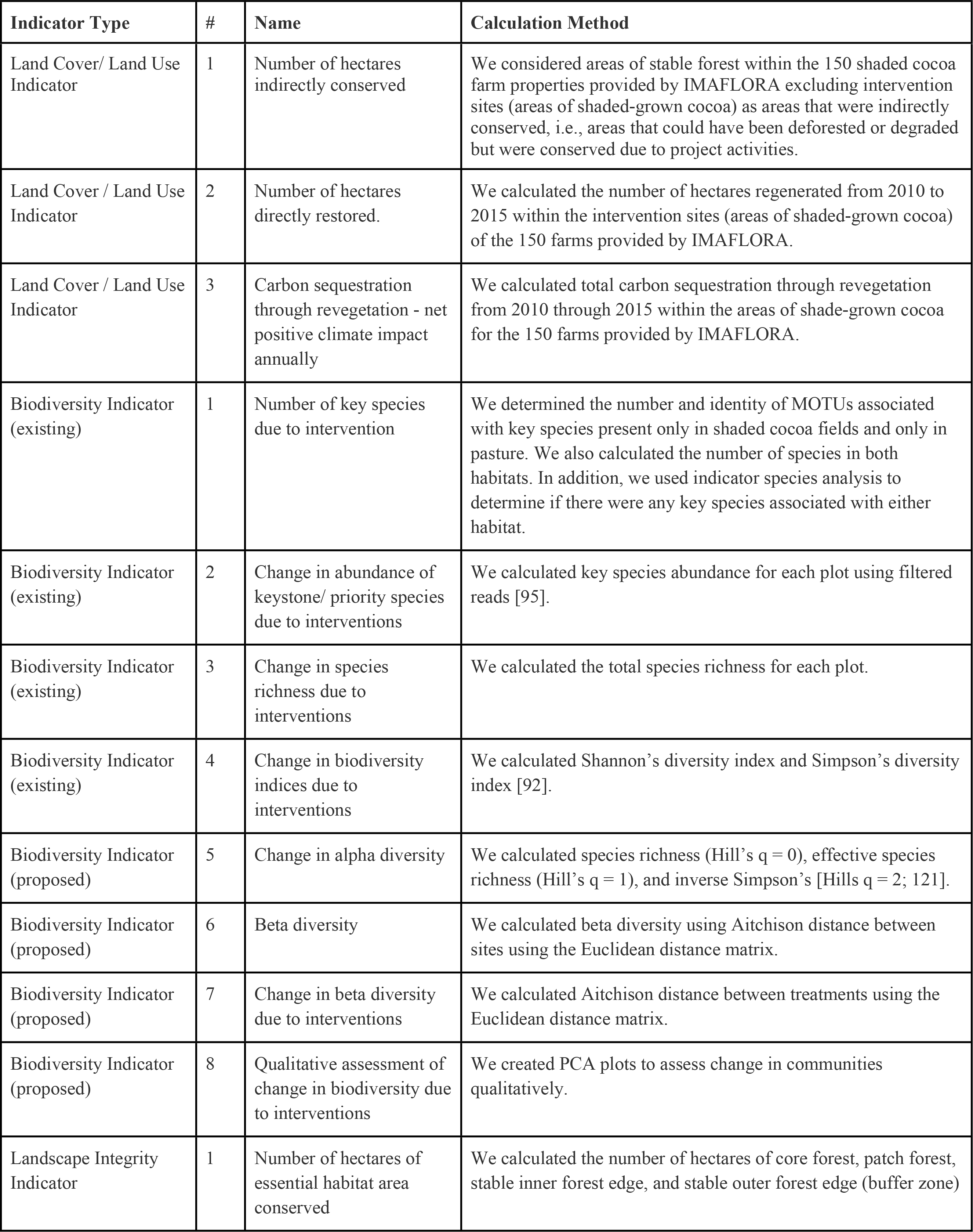
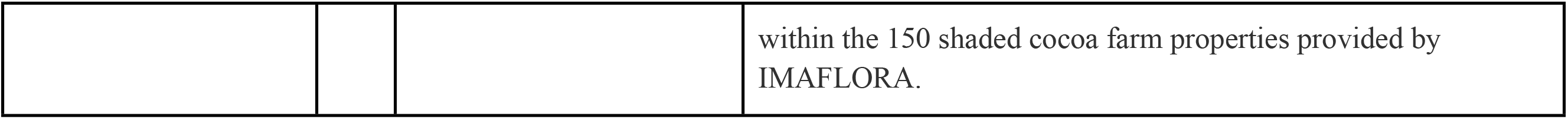
Overview of all indicators calculated for the TerraBio implementation.

In addition to these indicators derived from existing sources, we proposed a number of new indicators based on the literature [24–25, 119]. These proposed biodiversity indicators included: Alpha diversity; Beta diversity; Change in beta diversity due to the intervention; and a qualitative assessment of biodiversity change due to the intervention [25–26, 120–121].

## RESULTS

### MAPPING RESULTS

According to the final disturbance mapping results, 183 hectares were degraded, and 358 hectares were deforested within the 150 IMAFLORA properties (11,546 ha) between 2010-2015. Overall, producer’s and user’s accuracies (± 95% confidence interval) per class from the disturbances classified map (Figure 3) are summarized in Table 2. As expected, the stable classes (stable forest and non-forest) had higher user’s and producer’s accuracies and lower uncertainties compared to the dynamic classes (degradation and deforestation). Nevertheless, omission and commission errors happened within and across these two groups of classes (stable and dynamic classes). For example, in the dynamic classes (degradation and deforestation): most incorrectly classified pixels in the degradation class were omitted in the deforestation class and committed to the stable forest class, and both omission and commission errors in the deforestation class happened in the degradation class. The reasons behind these errors are explained in the discussion section.

**Figure 3:**
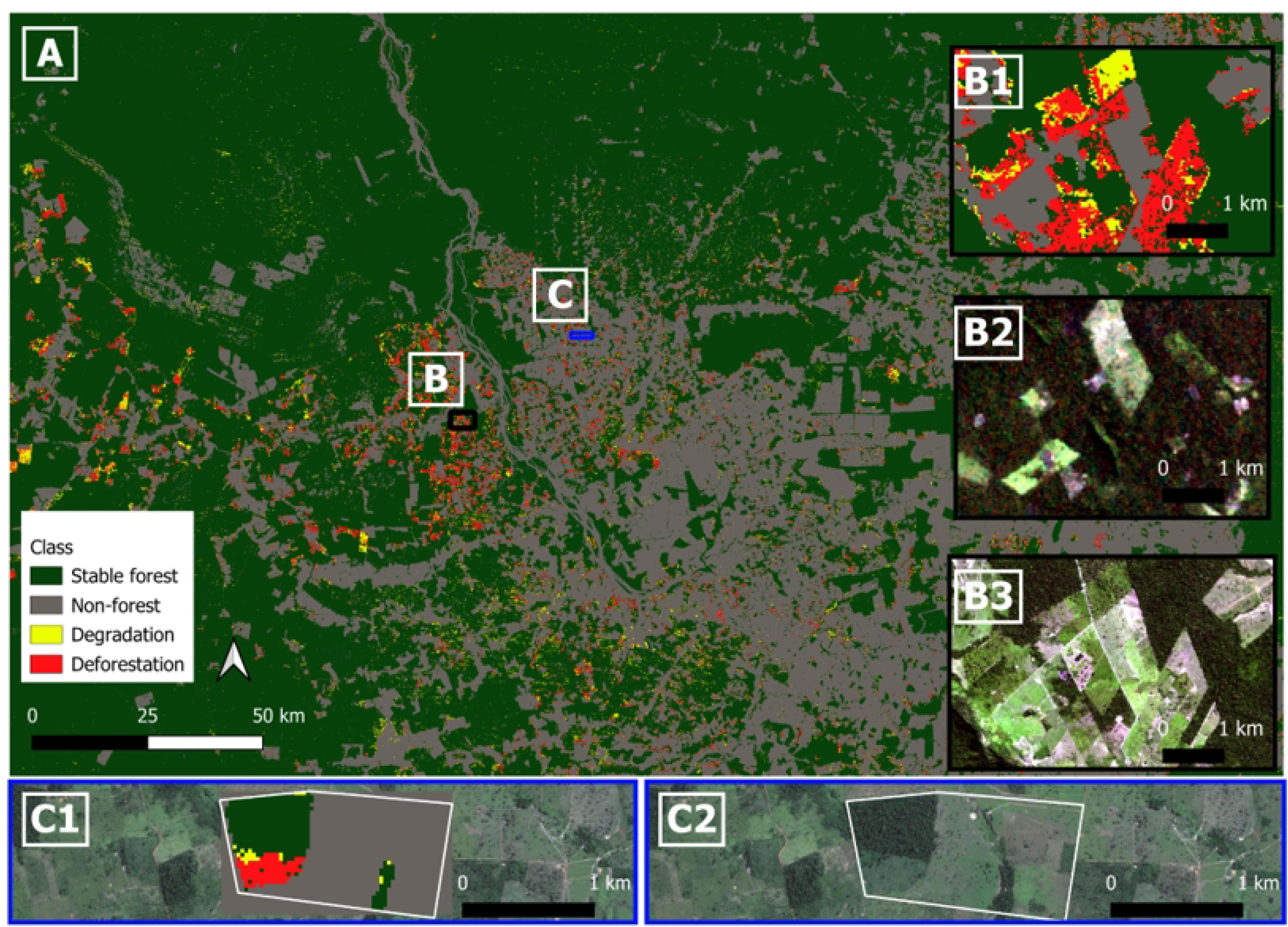
(A) Disturbance classification 2010-2015. Inset maps (B: B1, B2, B3) are shown with black outline and inset maps (C: C1, C2) are shown with blue outline on the main map. (B1) Inset map of disturbance classification 2010-2015 over a particular area to outline classification vs. RGB images. (B2) Inset RGB image pre-study period from Landsat 5 (July 30, 2009). (B3) Inset RGB image post-study period from Sentinel-2 (June 26, 2016). (C1, C2) Example of disturbances within one of the properties (degradation in yellow, deforestation in red, stable forest in dark green, and non-forest in gray). Note that the satellite image in (C1, C2) is from Google and does not have a timestamp associated with it.

**Table 2:** Accuracy assessment of the disturbances map.

| Accuracies (%) | Degradation | Deforestation | Stable forest | Non-forest |
| --- | --- | --- | --- | --- |
| User's accuracy | 40.5 ± 10.2 | 60.5 ± 9.1 | 89.0 ± 6.2 | 85.0 ± 5.4 |
| Producer's accuracy | 6.8 ± 3.4 | 30.5 ± 16.2 | 95.9 ± 1.8 | 97.0 ± 4.0 |
| Overall accuracy | 86.5 ± 4.3 | | | |

Overall, producer’s and user’s accuracies (± 95% confidence interval) from the regeneration classified map (Figure 4) are summarized in Table 3. As expected, our approach performed better at identifying where regeneration did not occur compared to where regeneration did occur. The regeneration class presented low user’s and producer’s accuracies and higher levels of uncertainties. The biggest error source was regeneration pixels being committed to the non-regeneration class, i.e., an overestimation of areas where regeneration occurred.

**Figure 4:**
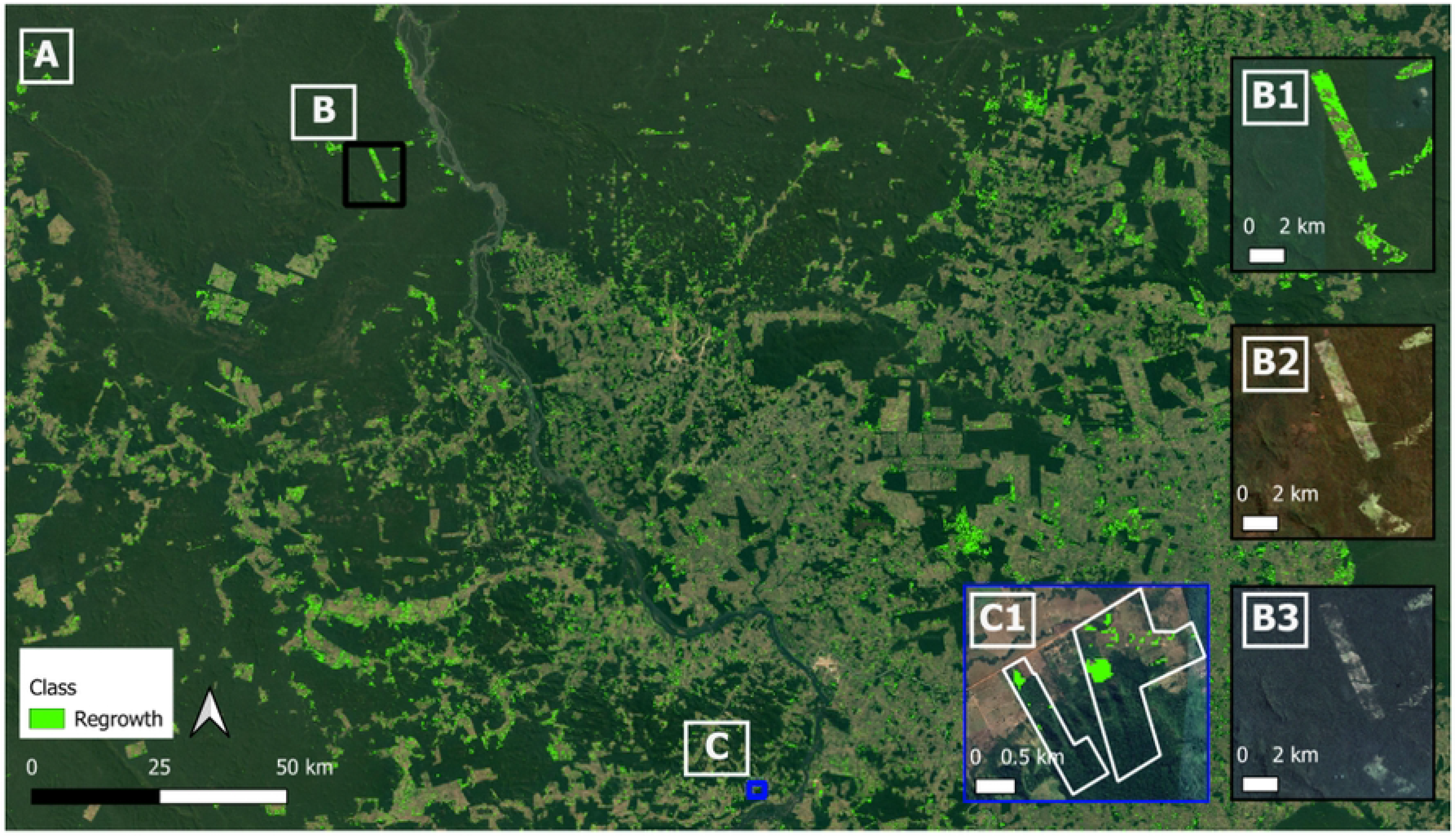
(A) regeneration map output 2010-2015. Inset maps (B: B1, B2, B3) are shown with black outline and inset maps (C: C1) are shown with blue outline on the main map. (B1) Inset map of regeneration output 2010-2015 over a particular area to outline regeneration areas vs. RGB images. (B2) Inset RGB median image pre-study period from Landsat 5 (July-August 2011). (B3) Inset RGB image post-study period from Planet NICFI mosaic (July -November 2016). (C1) Example of regeneration areas (in green) within two of the properties.

**Table 3:** Accuracy assessment of the regeneration map.

| **Accuracies (%)** | **Non-regeneration** | **Regeneration** |
| --- | --- | --- |
| **User's accuracy** | 98.3 ± 1.7 | 34.1 ± 9.1 |
| **Producer's accuracy** | 97.6 ± 0.3 | 43.0 ± 25.3 |
| **Overall accuracy** | 96.0 ± 1.6 |  |

### eDNA RESULTS

Overall, five cocoa sites and five pasture sites, in addition to one forest site, were surveyed by the field team. While the paired sampling design was not maintained, all but one of the fields sampled came from cluster 1 so the reduction in exogenous variation should be maintained (S3 Table).

Upon the initial bioinformatics filtering steps, a yield of 14,387,489 sequencing reads was obtained from the two sequencing runs, with 10,775,915 reads for the invertebrates dataset (5,598,018 Gillet, and 5,177,897 Zeale) and 3,611,574 for vertebrates. After applying all taxonomic filtering steps and retrieving only target MOTUs, the number of reads retained were 2,665,095 reads for arthropods including both Gillet and Zeale primer sets, and 38,806 reads for vertebrates after the removal of human and domestic animal reads. For the vertebrates dataset, a very high proportion of human reads was detected (>75% of total reads, from 6.9 million mammal reads, 5.2 million belonged to humans), with 14 MOTUs from 10 unique families identified when considering only the target wild mammalian taxa. For arthropods, 290 MOTU from 27 unique identified families were retained after the removal of MOTUs that were not assigned any taxonomic information. Due to the low amount of reads and MOTU returned for vertebrates, further analyses focused on the arthropod dataset (S3 File).

### INDICATOR RESULTS

**Land Cover/ Land Use Indicator 1: Number of hectares indirectly conserved.** We found 2,871 hectares within the 150 IMAFLORA properties (excluding intervention sites) were indirectly conserved between 2010-2015.

**Land Cover/ Land Use Indicator 2: Number of hectares directly restored.** We found 471 hectares within the intervention sites (areas of shaded-grown cocoa) of the 150 farms had regeneration between 2010-2015.

**Land Cover / Land Use Indicator 3: Carbon sequestration through revegetation - net positive climate impact annually.** We found carbon sequestration through revegetation in shade-grown cocoa systems in the intervention farms of 44,300 Mg C (8,860 Mg C/yr) between 2010 and 2015.

**Biodiversity Indicator 1: Number of key species due to intervention.** Overall, we found a total of 19 key MOTUs, including 11 in Hymenoptera and 8 in Lepidoptera. Most Hymenoptera belonged to the family Formicidae (ants), while Lepidoptera belonged to families Saturniidae and Crambidae, among others. Only one group of butterflies was detected (*Hermeuptychia hermes* [Fabricius], or the Hermes satyr).

One member of Hymenoptera was found in both cocoa fields and pasture. Five MOTU were found only in cocoa fields, including *Labidus* sp. (army ants), *Solenopsis* sp. (fire ants), *Hileithia* sp. (moth), a member of family Platygastridae and a member of Hymenoptera for which further identification was not possible. Thirteen MOTU were found only in pastures, including *Crematogaster abstinens* [Forel] and two other *Crematogaster* species, *Solenopsis geminata* [Fabricius] and another *Solenopsis* species*, Argyria* sp*., Heliura* sp., *Hermeuptychia hermes*, *Hylesia* sp. *Clepsis* sp., and a member of Hymenoptera for which further identification was not possible. No Hymenoptera or Lepidoptera were detected in forests. No species were consistently found to be indicator species.

**Biodiversity Indicator 2: Change in abundance of keystone/ priority species due to interventions.** As this pilot only included one time period, we compared the intervention to the counterfactual for this one time period. We found that there was also no significant difference in key species abundance when comparing cocoa fields (intervention) and pasture (counterfactuals; Pr(>Chisq) = 0.2648).

**Biodiversity Indicator 3: Change in species richness due to interventions.** The mean species richness in cocoa fields was 17.9 species (SD = 11.0), while on pastures mean species richness was 18.2 (SD = 9.4). When comparing total species richness between cocoa fields and pasture, we found there was no significant difference in MOTU richness (Pr(>Chisq) = 0.978; Figure 5).

**Figure 5:**
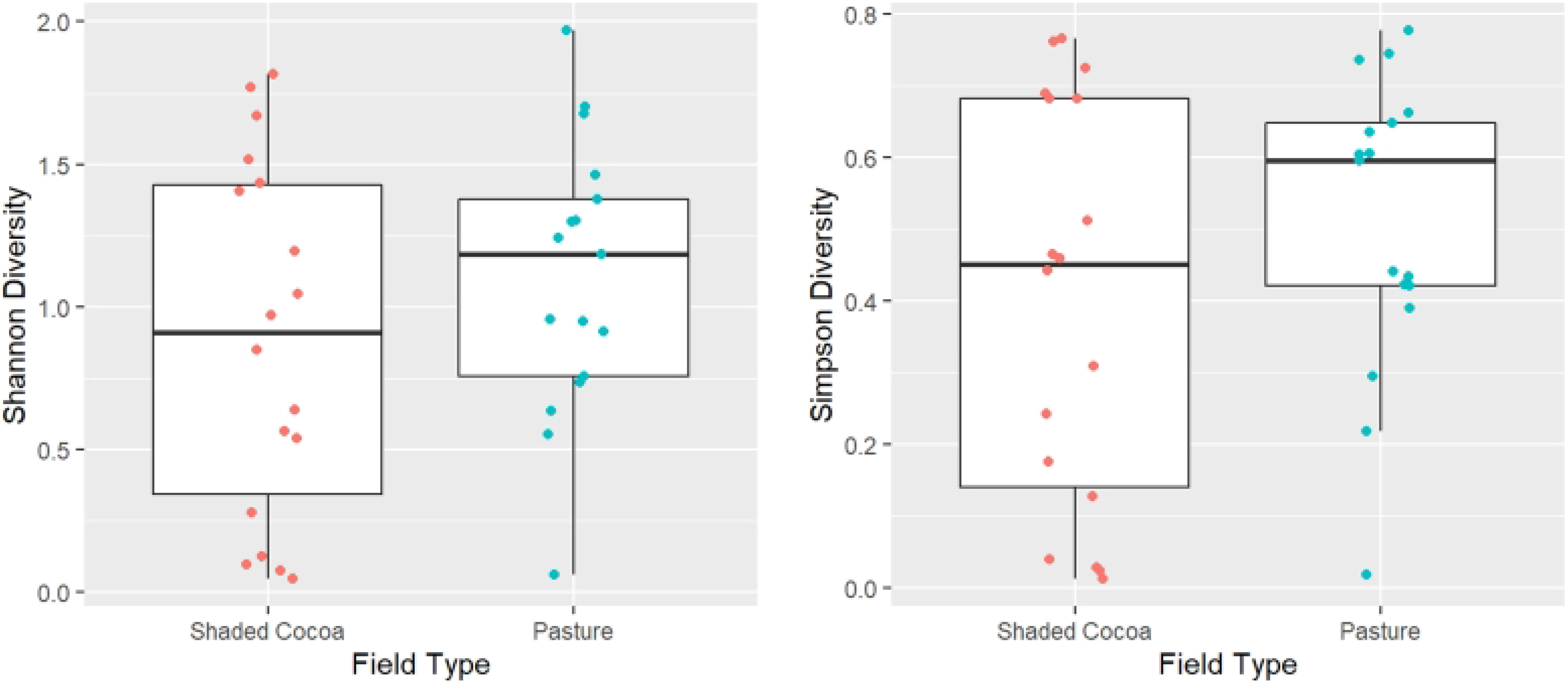
Species richness by field type. Each dot represents a sample point in shaded cocoa fields (red) or pasture fields (blue). Shannon diversity (left) and Simpson’s (right) by field type.

**Biodiversity Indicator 4: Change in biodiversity indices due to interventions.** When comparing diversity indices between shaded cocoa fields (intervention) and pasture (counterfactuals), we found that there were no significant differences for Shannon diversity (Pr(>Chisq) = 0.3639) or for Simpson’s diversity (Pr(>Chisq) = 0.2964; Figure 5).

**Biodiversity Indicator 5: Alpha diversity.** When comparing diversity indices between cocoa fields (intervention) and pasture (counterfactuals), we found that there were no significant differences for species richness (Hill’s q = 0; previous section), for effective species richness (Hill’s q = 1; Pr(>Chisq) = 0.4824), and for inverse Simpson’s (Hill’s q = 2; Pr(>Chisq) = 0.5243; Figure 6).

**Figure 6:**
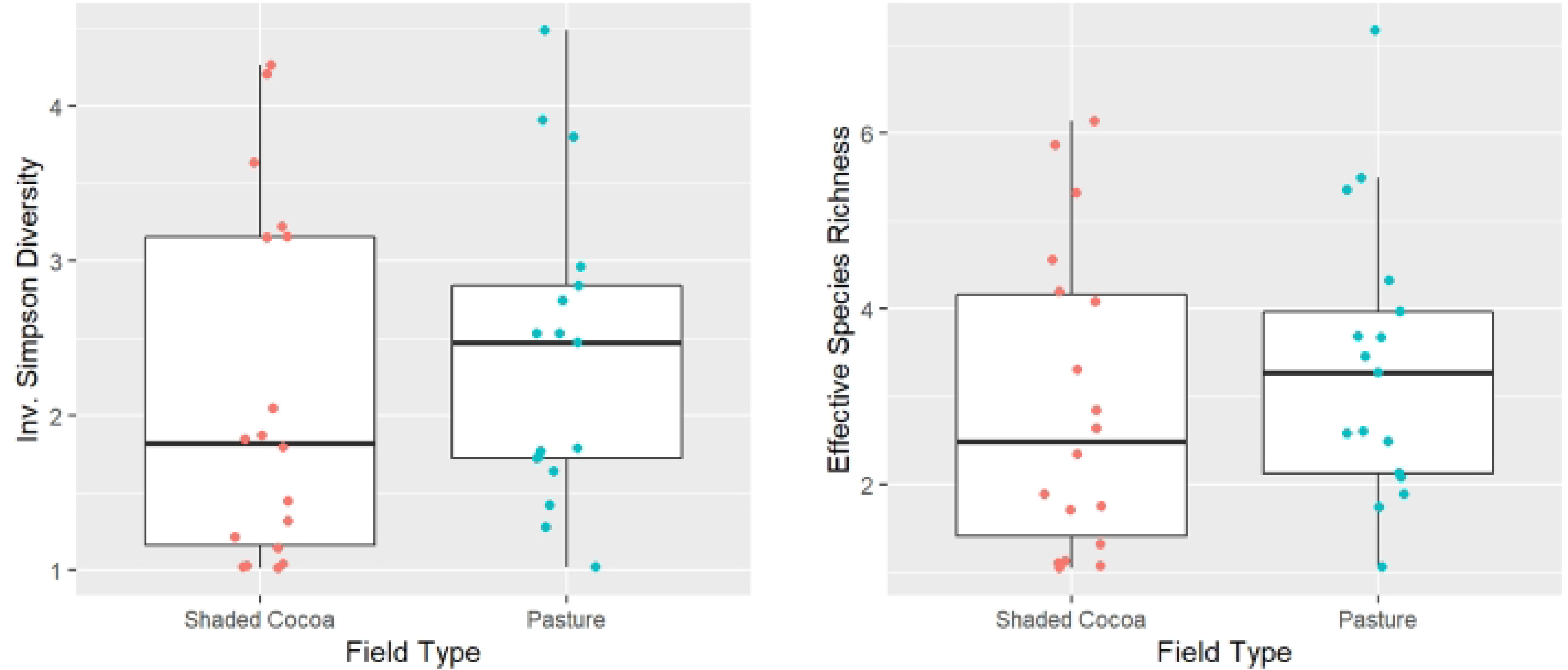
Effective species richness (L) and Inverse Simpson Diversity (R) by field type. Each dot represents a sample point in shaded cocoa fields (red) or pasture fields (blue).

**Biodiversity Indicator 6: Beta diversity.** Pairwise distances between all sampling sites ranged from 10.3 and 35.1, where 0 represents no dissimilarity, and larger distances indicate increasing dissimilarity (Figure 7). The highest dissimilarities were observed between the Cocoa 1 field and the Pasture fields.

**Figure 7:**
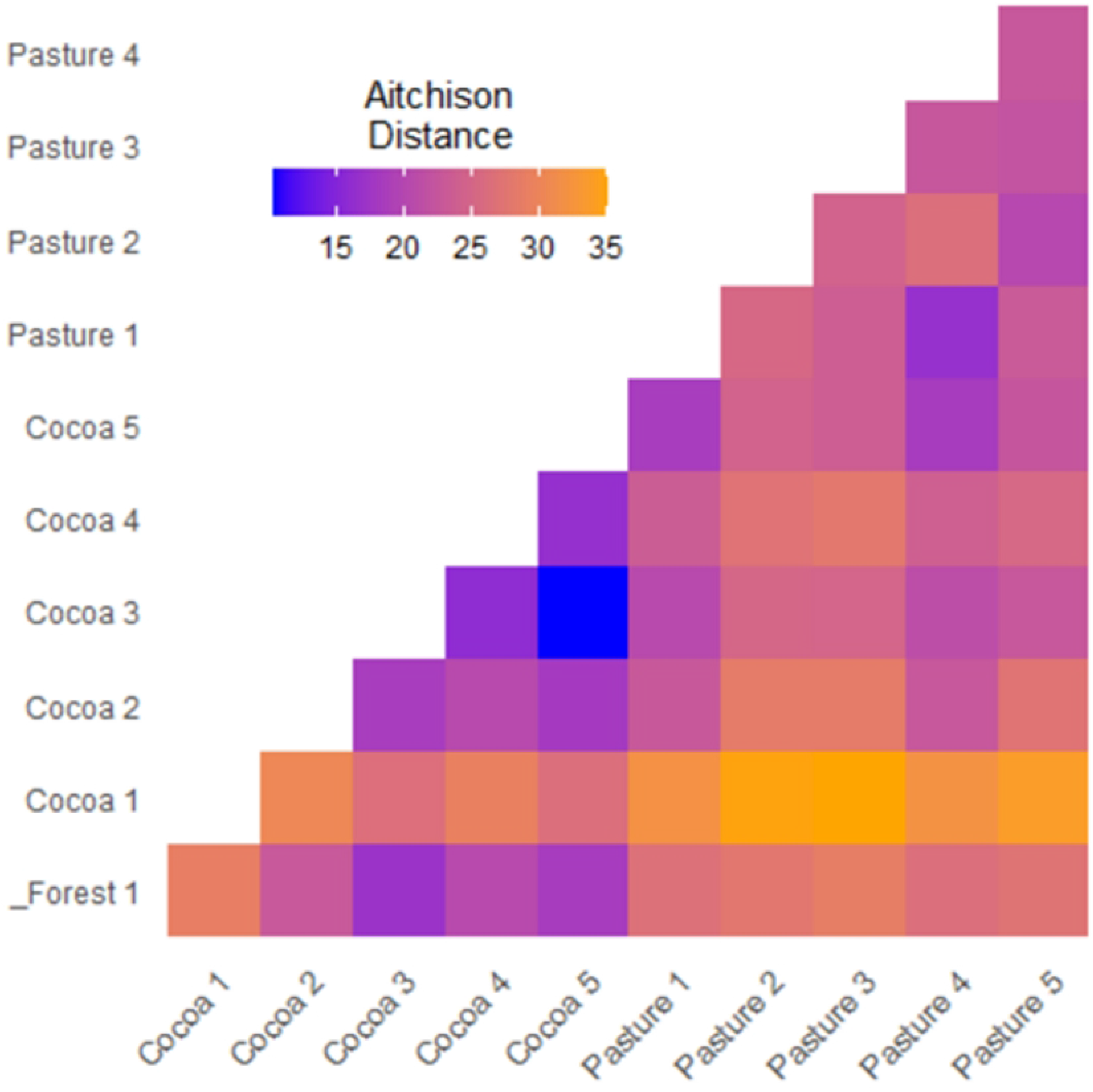
Beta diversity (Aitchison distance) between each site.

**Biodiversity Indicator 7: Change in beta diversity due to the intervention.** The Aitchison distance between shaded cocoa (intervention) and pasture (BAU) was about equal to the distance between shaded cocoa (intervention) and forest (control; 16.6 and 16.8 respectively), but much smaller than the distance between pasture (BAU) and forest (control; 23.2), though this difference was not significant [122; Figure 8].

**Figure 8:**
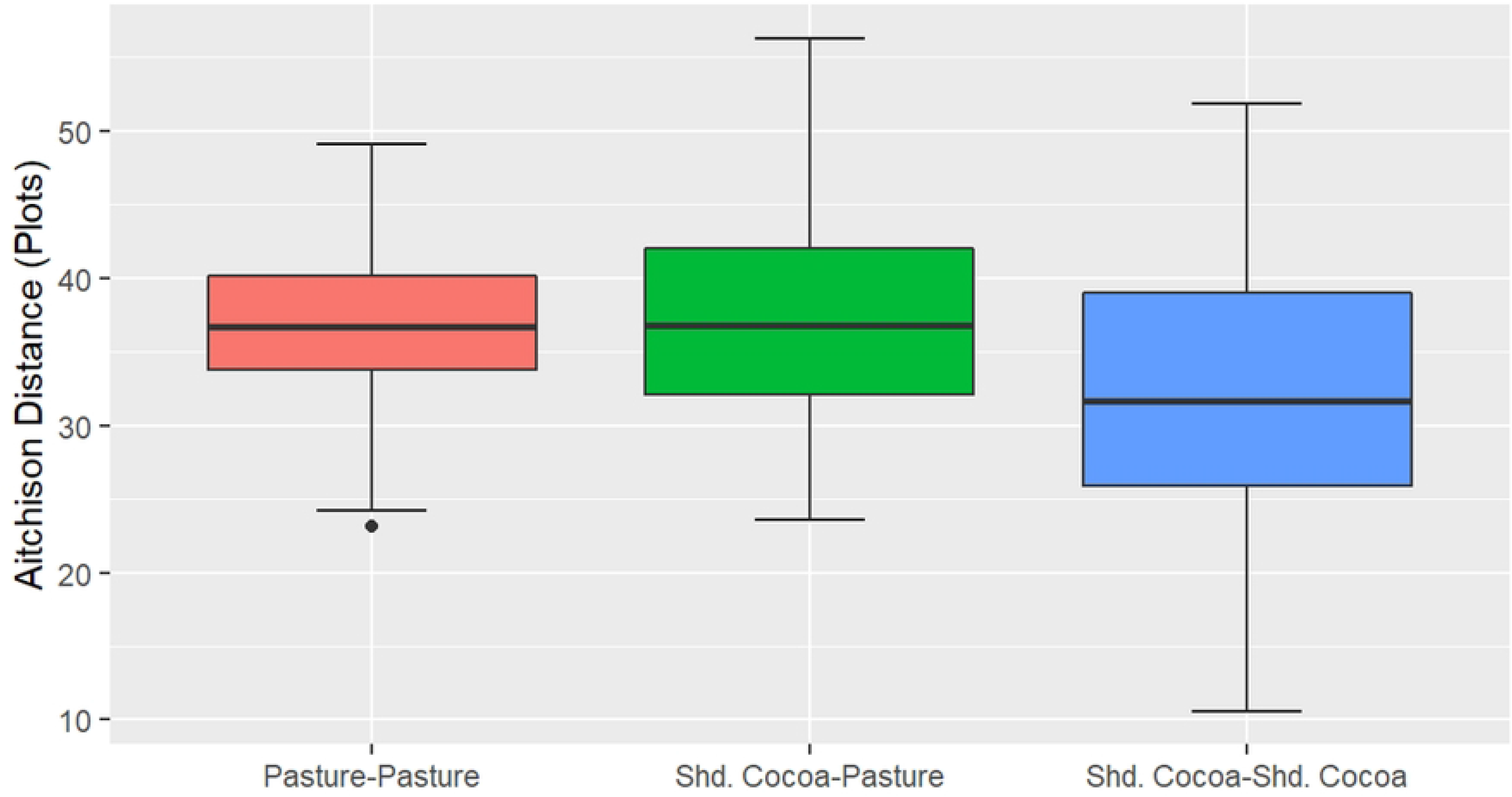
Aitchison distances within treatments and between treatments.

**Biodiversity Indicator 8: Qualitative assessment of change in biodiversity due to the intervention.** PCA graphs of plots and sites revealed that cocoa and forest sites were co-located in site-species space, while pastures were strongly separated along the first axis (Figure 9). This agrees with our findings in Biodiversity Indicator 7.

**Figure 9:**
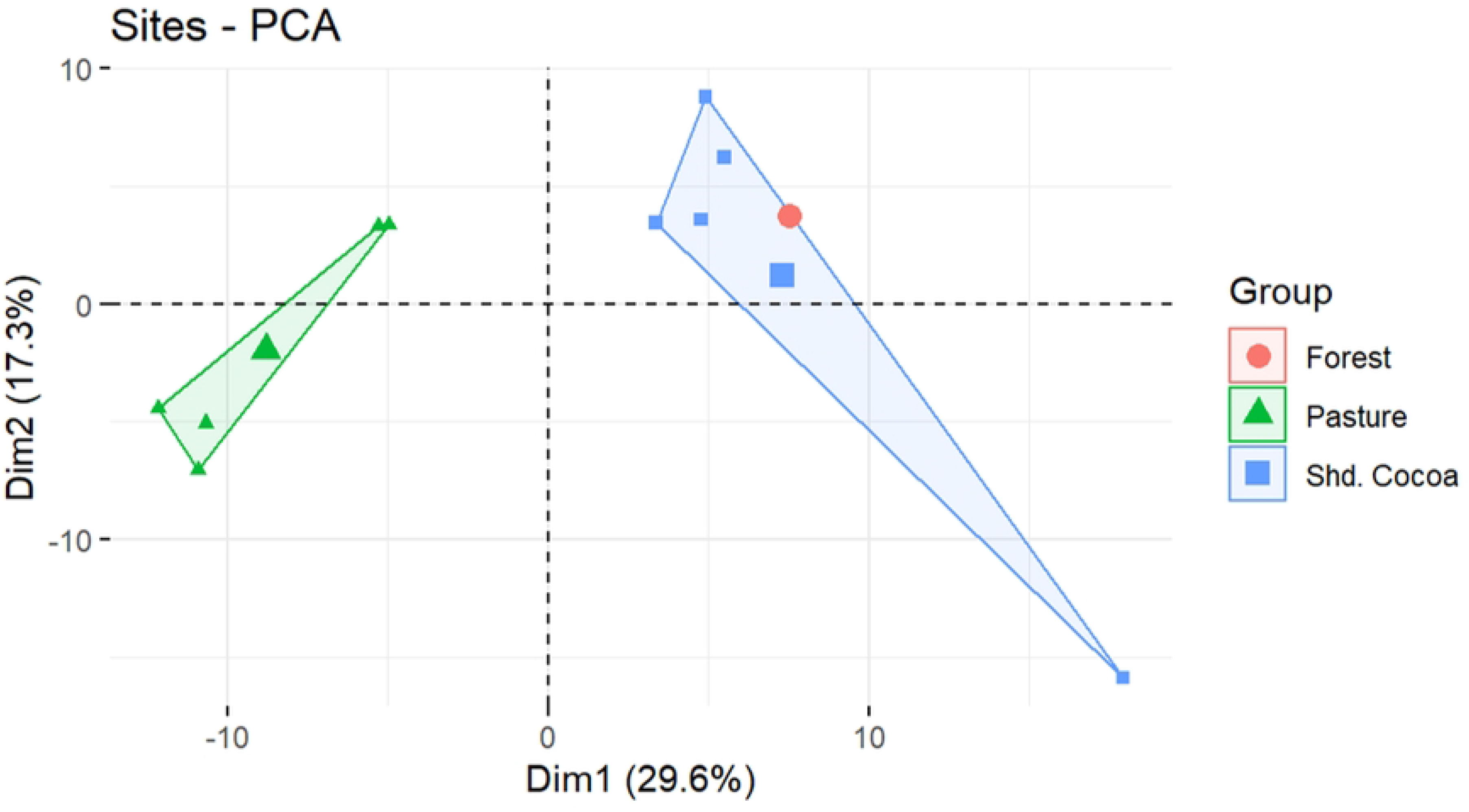
Arthropod PCA for arthropods found in shaded cocoa, forest, and pasture sites.

**Landscape Integrity Indicator 1: No. of hectares of essential habitat area conserved.** Essential habitat areas calculated using MSPA within all farm properties with shaded cocoa included 93 ha of core forest, 114 ha of patch forest, 82 ha of stable inner forest edge, and 1,000 ha of stable outer forest edge (buffer habitat; S2 Table). Thus, a total of 289 ha can be considered critical habitat within the intervention farms, and an additional 1,000 ha is stable buffer habitat.

## DISCUSSION

### PILOTING THE TERRABIO FRAMEWORK

The pilot application of TerraBio, a coupled eDNA and remote sensing environmental monitoring approach, demonstrated the potential for such systems in monitoring sustainable agriculture such as the shaded cocoa fields implemented by IMAFLROA. Overall, our remote sensing analysis suggests that the shaded cocoa established by IMAFLORA helped revegetate over 400 hectares, and the eDNA analysis suggests that the community composition of arthropods in shaded cocoa is closer to second growth forests than that of pastures.

Based on the indicators, the broader impacts of shaded cocoa in the study area are most likely increased canopy cover, increased carbon sequestration, and more ‘forest-like’ habitat availability for arthropods. Land Cover/Land Use Indicators 1 and 2 both found support for the shaded cocoa indicators directly and indirectly restoring forest cover. Land Cover/Land Use Indicator 3 similarly found a net gain due to carbon sequestration by the shade canopy and cocoa in the intervention farms. While the results from the initially provided biodiversity indicators and vertebrates were inconclusive, the proposed biodiversity indicators found that arthropod communities in shaded cocoa fields were closer to forests than arthropod communities in pastures were to forests. This suggests that the habitat available to arthropods in shaded cocoa was more ‘forest-like’ than the pastures due to the interventions, in agreement with Landscape Integrity Indicator 1.

Importantly, our results agree with previous studies on the impacts of shaded cocoa. In Ethiopia, researchers found that shade coffee certification increased the probability of forest conservation by 19.3% [123] and that indirect forest conservation was also observed within 100m of the project areas [124]. Similarly, the contribution of shaded cocoa to biodiversity conservation viewed through the lens of retaining forest-like communities echoes decades of previous research [125–128]. Contributions of shaded cocoa systems to landscape connectivity are also well supported [127, 129], though systems emphasizing native trees are likely more successful than those using bananas and *Erythrina fusca* [130]. In Brazil’s Atlantic Forest, traditional agroforests where cacao is planted under thinned native forests called *cabrucas,* have greater diversity of tree species, including forest specialist tree species, and while they are not substitutes for undisturbed forest they do have a critical role in biodiversity conservation [131].

### LEARNING FROM THE PILOT IMPLEMENTATION

Implementing the pilot allowed us to both improve TerraBio and provide guidance for other coupled monitoring approaches for the future. In general, we found multiple aspects of our approach worked well, including robust field sampling using random samples, using the management unit as the unit of analysis, and benchmarking against a control and/or business as usual land use.

### LESSONS FOR REMOTE SENSING

With the remote sensing component of the coupled methodology, we encountered some challenges specific to the complex agroecosystems of the Brazilian Amazon, highlighting the need for careful integration of region-specific remote sensing knowledge in developing coupled monitoring approaches. These were detected during the accuracy assessment, highlighting the importance of this step.

First, to calculate the Land Cover/Land Use and Landscape Integrity indicators, we created two map products. While the methods used to create these map products have significant support in the literature [e.g., 51, 133-134], we found that applying them in this specific context had wide margins of error and some unexpected results that provided an opportunity to learn and improve upon these methods. For example, our disturbance maps, which were used for LCLU Indicator 2, showed omission errors in the stable forest class. In accordance with [134], many of the omission errors associated with this class were derived from the presence of deciduous tree species (“caducifólias’’, in Portuguese) in this region, which show a seasonal leaf color change, leaf-off pattern (Figure 10), and changes in NDFI values. Therefore, the algorithms assume a disturbance event in forested lands, classifying most of these pixels as degradation, which explains the low accuracy obtained for these classes. Both CODED and LandTrendr should be able to capture seasonal variations of forests with varying crown covers, and parametrization for local conditions can mitigate this issue [46–48].

**Figure 10:**
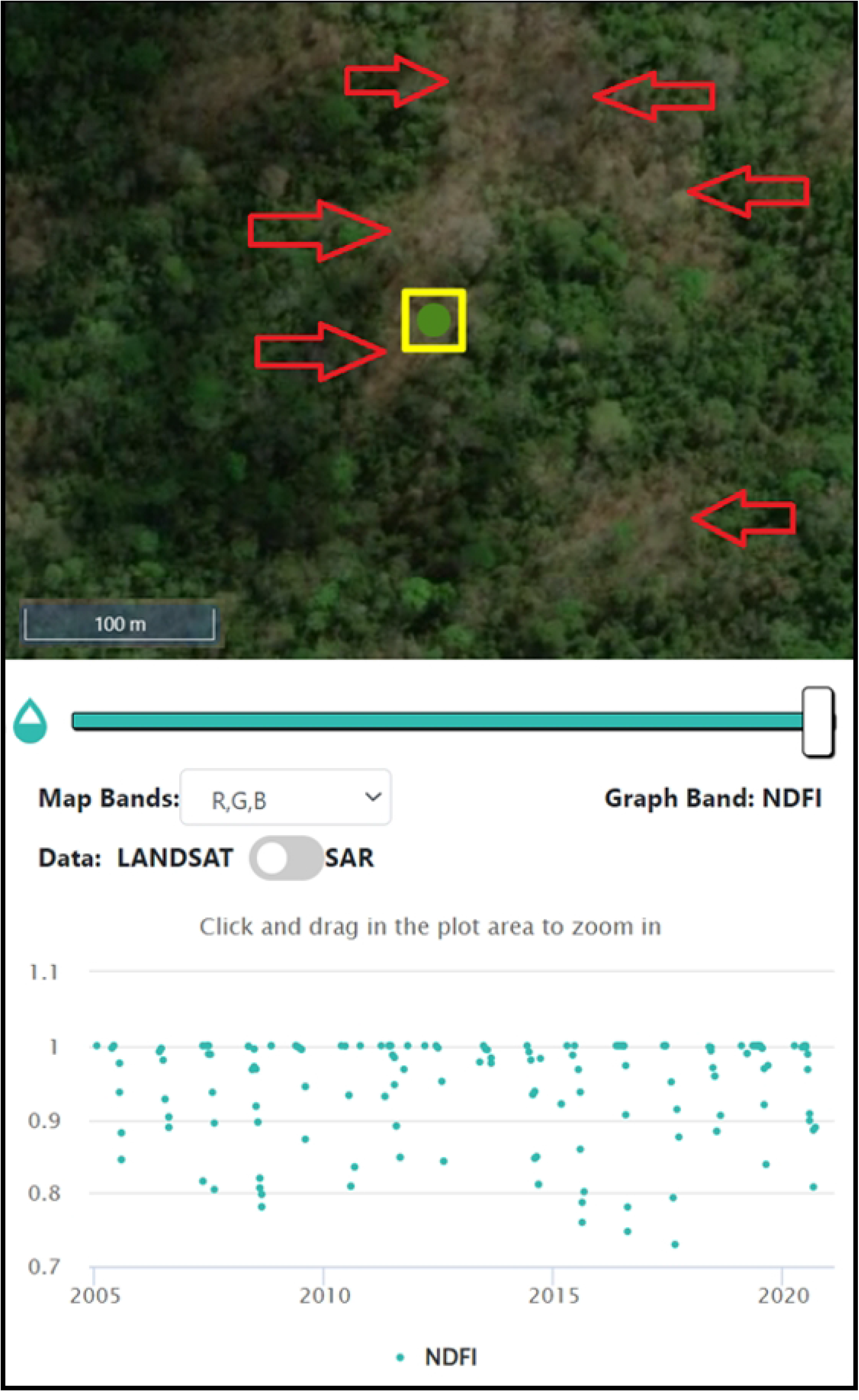
Plot in the CEO disturbances validation project. This pixel was misclassified as Degradation. NDFI time series show a seasonal pattern with lower NDFI values around August of each year. It is important to note that the MapBiomas classification maps these areas as “Savannic Forest Formations”.

Further, it is important to fully understand the forest conversion process and pastureland management practices in the region, as this greatly influences the interpretation itself of disturbance samples for validation. Many of the samples analyzed represented patches of degraded forest cover or “dirty pastures”. Some of the degraded patches presented recurrent regeneration and disturbance patterns, visible in the NDFI time series due to fragmentation or border effect (Figure 11). This is sometimes not captured in the visual interpretation analysis, where just one change event is recorded, which yields omission or commission errors for the change classes (e.g., regeneration and degradation, deforestation and degradation). For future applications, it will be important to separate the regeneration output validation from the disturbance output validation to refine confidence intervals.

**Figure 11:**
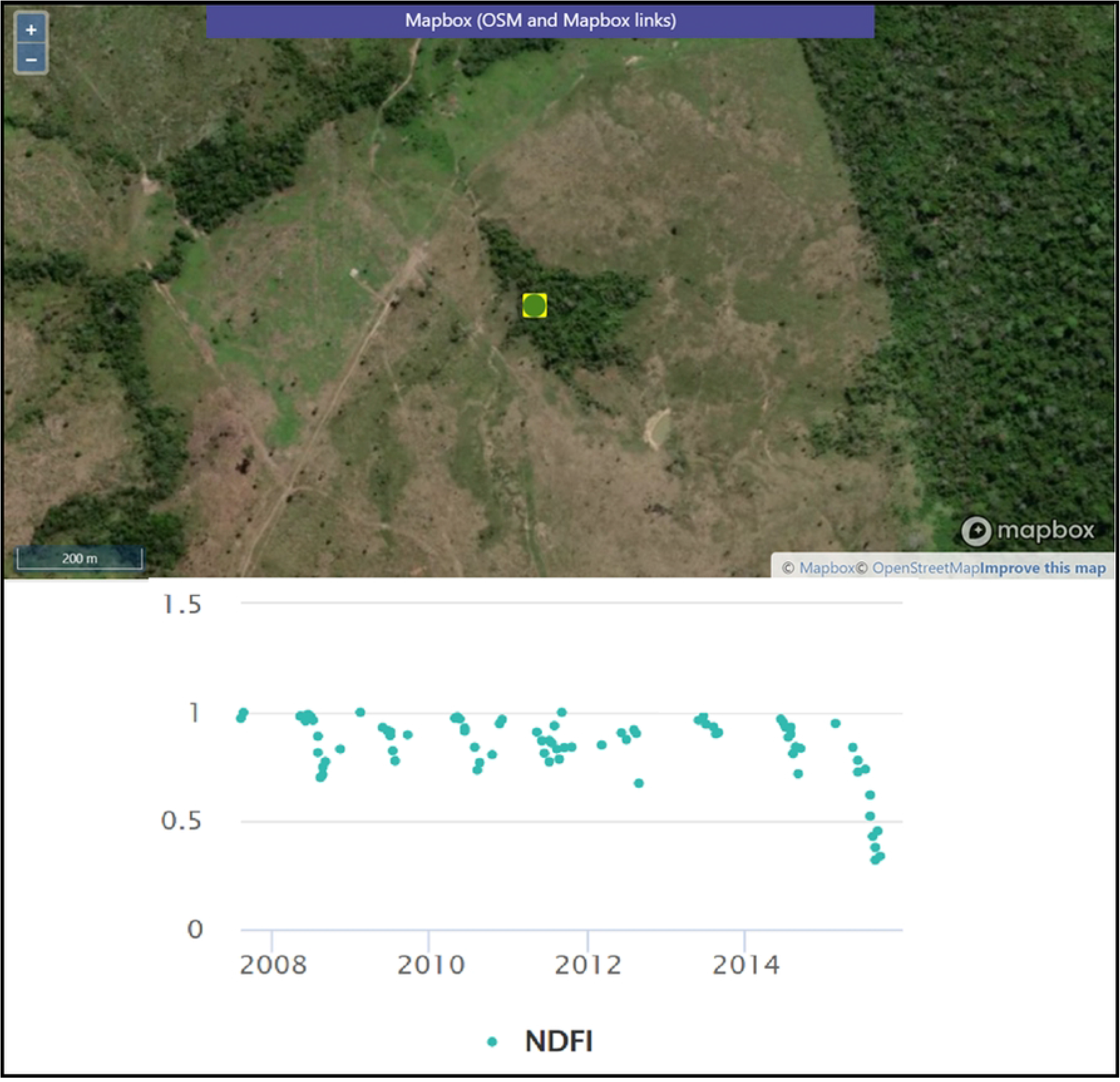
Example plot of the CEO change validation project showing a fragmented/degraded forested patch and its variations on the NDFI time series.

A better understanding of the degradation process and its relationship with deforestation is vital for the decision-making process of interpreting validation samples. Many times, the disturbances happen gradually, in phases, until they reach the final clear-cut stage of deforestation. The relationship between degradation and deforestation may vary significantly across the different land tenures. The same happens with the regeneration process, which will be characterized by secondary forests with different stages depending on how many years of recovery we are seeing. The interpretation of what is happening on the ground is not always clear and straightforward, especially when the interpretation is being made through satellite imagery (Figure 12). In a span of five years, we may observe two or three different events (e.g., selective logging followed by fire and then regeneration). Without the availability of Planet NICFI data prior to 2015, the visual inspection of these events through Landsat imagery and NDFI time series interpretation was challenging, which could have yielded validation errors.

**Figure 12:**
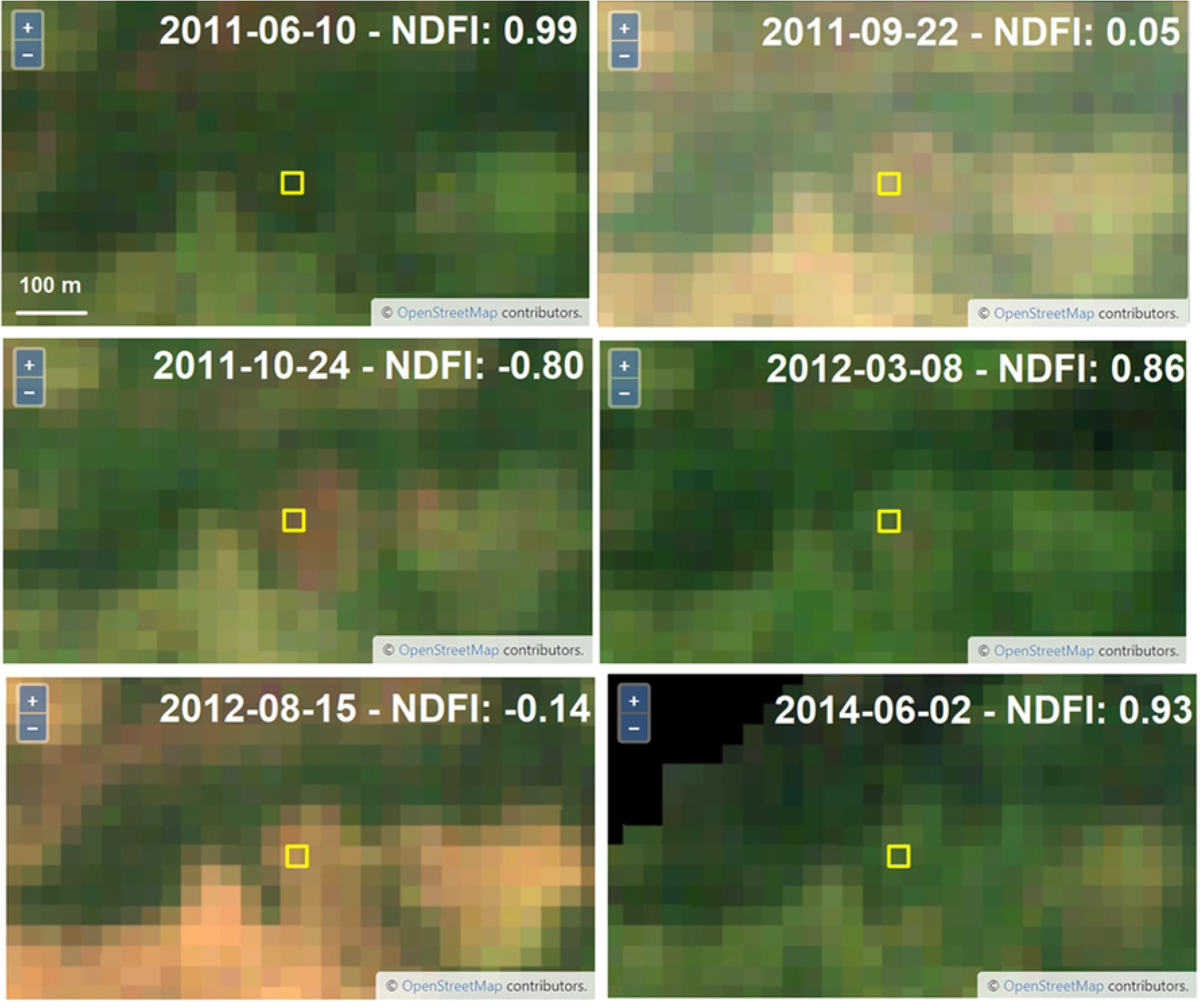
Example plot of the CEO change validation project showing a plot that was classified as regrowth by the algorithm, but the interpreter classified it as a single degradation event. We note some regreening between the dates and the variation in NDFI values. Although not entirely clear, the interpreter assumed selective logging followed by a fire event in 2011. Another fire event seems to have happened in 2012. It is not clear that by the end of 2014 the area was already-established pastureland.

Finally, our estimates of carbon sequestration due to revegetation do not align well with those that have been estimated previously. For example, potential sequestration for smallholder agroforestry systems like shaded cocoa has been estimated at 1.5 to 3.5 Mg per year per hectare [135]. This rate suggests that we should expect approximately 6,000 Mg C sequestration due to revegetation over the study period based on our regeneration maps. Our overestimate is caused by the approach’s treatment of regeneration as entirely occurring within the study time period, which is unrealistic. For future applications of TerraBio, we plan to adapt the carbon calculation method by [136]. We will use forest stand age calculated by the LandTrendr algorithm to create more accurate estimates of carbon sequestration through revegetation by more realistically accounting for the rate of growth and thus rate of sequestration.

We also recognize similar limitations with the MSPA calculations by using the Global Forest Change dataset. We plan to use the LandTrendr product and simplify the MSPA classes. By applying the existing LandTrendr product, and therefore, more local-based information, to both methods we expect the results to have higher accuracy, which can be supported by confidence intervals for the case of the carbon estimates.

### LESSONS FOR MONITORING WITH eDNA

The success of using arthropod communities as indicators focused more on community composition points to some lessons learned for future coupled biodiversity monitoring approaches and implementations of TerraBio. The arthropod datasets did not encounter the same issues with high numbers of human reads and domesticated species, likely associated with the anthropogenic land uses [137–138]. Combined with their ecological importance, arthropods are thus a promising taxonomic group to use as a target moving forward [97, 139]. Both Gillet and Zeale primers succeeded in capturing different parts of the Arthropoda phylum, however most reads obtained belonged to non-target taxa, and for future studies, we recommend that both primer sets be used if both Hymenoptera and Lepidoptera groups are targeted taxa.

From the reads attributed to Arthopoda, a significant fraction of detected MOTUs could not be successfully assigned at the taxonomic rank required for the downstream analyses. The lack of resolution at short fragments associated with eDNA monitoring and sparse or incomplete reference databases is a well-known issue in understudied regions such as the Neotropics [140]. For the “key species” indicators requiring identification to species, many key arthropod species were present in only one plot. Thus, when monitoring these indicators over time, as TerraBio plans to do, researchers must either use an outdated database or constantly update previous years’ results in order to separate the effect of improving databases from any real biological change. Further, due to the fundamental character of eDNA data, some traditional indicators such as species abundance calculated with eDNA data can be misleading [119].

This points to one key benefit of using indicators focused on overall community composition over those focused on species specific (e.g., keystone, rare, endangered, or endemic species) used elsewhere. The initial set of indicators selected for use in the pilot implementation of TerraBio were derived from existing indicators based on traditional field ecology methods where organisms are directly observed. However, eDNA represents a fundamental shift in how ecological communities are measured. Any taxonomy-based indicator requires accurate databases linking genomic sequences with their taxonomic identity [19, 25]. New approaches to ecological indicator selection can better leverage eDNA data to monitor communities using taxonomy-free approaches [24–25, 119].

Our proposed indicators, chosen specifically to take advantage of eDNA data, were much more successful at identifying differences between communities found in the intervention. For example, Biodiversity Indicators 2-4 found no significant differences between the shaded cocoa and pasture sites, however, the community focused Biodiversity Indicators 6-8 all found that cocoa fields were more similar to forest plots than the pasture sites. Using these community ecology-based taxonomy-free approaches to fully make use of eDNA data is an important lesson learned for coupled environmental monitoring approaches, including TerraBio. However, when implementing this approach to indicators, they must be clearly communicated to stakeholders accustomed to the outputs of traditional field ecology methods. Overall, eDNA is well suited to monitoring these projects, but successful integration in monitoring, reporting, and verification (MRV) standards will require balancing the best available science with reporting requirements.

We also encountered some challenges with our sampling protocols when implemented in remote areas by field partners and packaging scientific best practices in a way that is easily accessible and actionable. During data collection, our field team ran into multiple practical issues while collecting soil samples using the plot sample design. Having specific areas to collect from proved very time consuming, which when added to the already long transit times between field locations resulted in reduced data collection. The size and shape of the fields also limited the field team’s ability to change the location of plots if necessitated by field conditions. These challenges will become increasingly important when farm owners and community scientists are collecting data.

Thus, we suggest using a simplified larger volume sampling approach, and will be moving to this in future implementations of TerraBio. In particular, we are moving away from a plot-based approach with small amounts of soil collected to an approach collecting large volumes of soil from the entire site [1-2 litres; 141]. Recent research suggests that large soil volumes are likely needed to accurately capture community representation [e.g., 141-144]. This sampling approach will also be significantly easier for farmers and community scientists to implement, and early testing is promising.

Further, during our data analysis, we noticed that fewer reads were returned for cocoa fields than for pasture fields, with multiple plots missing data entirely or exhibiting very low species richness (i.e., number of detected species, S3 Table). Differences in soil moisture may have accounted for this result, as pastures were generally drier than shaded cocoa fields and forests. If the desiccant volume was not sufficient to handle higher soil moistures, then sample degradation may have occurred unevenly between the more shaded and thus wetter cocoa and forest samples and the drier pasture samples [138]. We will be testing higher volumes of desiccant and Longmire’s solution, as many of the sustainable agricultural projects occur in remote areas and tropical or sub-tropical climates [145–146]. Early results suggest this approach may help with high moisture soils, allowing for more accurate comparisons between land uses.

## CONCLUSION

Approaches that directly monitor land cover change and biodiversity on an annual basis by coupling remote sensing and environmental DNA (eDNA) can provide timely and relevant results for parties interested in the success of sustainable agricultural practices. Monitoring information collected from these approaches serves two essential functions: assessing the effectiveness of project-level management actions and approaches and facilitating ongoing learning about the circumstances in which different approaches outperform others.

In this pilot, we found that shaded cocoa projects implemented by IMAFLORA contributed to forest and biodiversity conservation in the Brazilian Amazon. The broader impacts of shaded cocoa projects in the study area, as revealed through the indicators, are most likely increased canopy cover, increased carbon sequestration, and more ‘forest-like’ habitat availability for arthropods. Following the successful pilot application of TerraBio to assess the effect of shaded cocoa interventions near São Félix do Xingu, in Pará, Brazil on biodiversity conservation, the approach will be expanded to other projects created through innovative funding mechanisms.

Implementing this pilot allowed us to provide guidance for coupled monitoring approaches for the future, including TerraBio. In general, we found multiple aspects of our coupled approach worked well. These included our sampling design using randomly selected farms and using the management unit as the unit of analysis. Benchmarking the intervention against a control and business as usual land use was a unique and cost-effective way to validate results on the project level while controlling for exogenous variables. Using remote sensing and eDNA data collection in tandem to calculate indicators also provided a more holistic view than either alone.

For our remote sensing analysis, we found that detecting forest disturbances and regeneration was challenging due to regional land management practices and vegetation characteristics, suggesting better algorithm parametrization to local conditions is needed to improve future accuracy. Understanding the degradation process and its relationship with deforestation is also vital for interpreting validation samples. Additionally, more realistic approaches to forest growth for carbon calculation analysis are needed. For our eDNA analysis, we suggest moving to straightforward sampling designs using high volume sampling and replication. We also found that taxonomy-free arthropod community focused indicators were more successful at illuminating ecologically holistic differences between intervention (shaded cocoa) and business as usual (pasture) scenarios.

## ACKNOWLEDGMENTS

We want to thank the farmers who allowed us to sample their fields for the eDNA work and Eric Bullock, for his expert advice on the use of his CODED algorithm in the Amazon.

## DECLARATIONS

### ETHICAL APPROVAL

No human or animal subjects were included in this research.

### AUTHOR CONTRIBUTIONS

**Conceptualization**: K.D., A.P.N., W.F., A.D., B.C., N.C., N.S., A.D.M., B.R., B.Z., D.S., and K.T.

**Data curation**: K.D., A.P.N., N.G.S., A.D.M., T.N.

**Formal analysis**: K.D., A.P.N., N.G.S., A.D.M., T.N., J.D., C.W., and K.T.

**Funding acquisition**: W.F., A.D., D.S., and K.T.

**Investigation**: K.D., A.P.N., W.F., A.D., B.C., N.C., V.C., G.A., C.L.P, J.T., S.C., O.R., N.G.S., A.D.M., I.J., C.W., B.Z., and K.T.

**Methodology**: K.D., A.P.N., W.F., A.D., N.S., A.D.M., B.Z., and K.T.

**Project administration**: W.F., A.D., C.W., D.S., and K.T.

**Resources**: W.F., A.D., D.S., and K.T.

**Supervision**: W.F., A.D., D.S., and K.T.

**Visualization**: K.D., A.P.N.

**Writing - original draft**: K.D., A.P.N.

**Writing - review and editing**: K.D., A.P.N., W.F., A.D., G.A., B.C., S.C., N.C., J.D., V.G., I.J., I.M.M., A.D.M., L.M., D.M.N., T.N., C.L.P., J.P., T.R.D., O.R., B.R., N.S., J.T., C.W., B.Z., K.T., and D.S.

## SUPPORTING INFORMATION

**S1 File. MPSA Definitions.** A detailed description of all MPSA definitions.

**S1 Table. Sample Sizes for Change Map Validation.** Sample size per class for validation efforts. The change maps (disturbance map and regeneration map) were combined into one map and the same sample dataset was used for accuracy assessment.

**S2 File. eDNA Soil Sampling Protocol.** Detailed protocol for eDNA soil sampling.

**S2 Table. Detailed MSPA Results.** Morphological Spatial Pattern Analysis of forest change dynamics from 2001 to 2019 in the study area.

**S3 Table. Biodiversity Results by Site.** Planned sampling and plots returning usable data after filtering.

**S3 File. Vertebrate Results.** Indicator results from the vertebrate primers.

## COMPETING INTERESTS

The authors have no relevant financial or non-financial interests to disclose.

## DATA AVAILABILITY

The datasets generated during and/or analyzed during the current study are available in the GitHub repository. Data will be published in the public repository at manuscript acceptance.

## Notes

### Competing Interest Statement

The authors have declared no competing interest.

